# Dengue transmission risk in a changing climate: Bangladesh could experience a longer dengue fever season in the future

**DOI:** 10.1101/2021.05.24.443300

**Authors:** Kishor K Paul, Ian Macadam, Donna Green, David G Regan, Richard T Gray

## Abstract

Our changing climate is already affecting the transmission of vector borne diseases such as dengue fever. This issue presents a significant public health concern for some nations, such as Bangladesh, which already experience regular seasonal outbreaks of dengue fever under present day conditions. To provide guidance for proactive public health planning to potentially mitigate future infections, we explore the impact of climate change on dengue infections by calculating the change in vectorial capacity of *Aedes aegypti* mosquito at a seasonal level for all regions in Bangladesh under two scenarios for future atmospheric greenhouse gas concentrations. For each of the four climate models used, and for both scenarios, our analysis reveals that the annual vectorial capacity remains at a level that would enable potential dengue epidemic transmission in all regions in Bangladesh. We found a slight decline in vectorial capacity in half of the regions examined during the last two decades of the 21^st^ Century for the lower-concentration scenario, with a pronounced decline in vectorial capacity in all geographic regions beginning in 2060 for the higher-concentration scenario. The likely reason is that in many regions, warming is leading to sub-optimal mosquito breeding temperatures. However, seasonal differences in vectorial capacity dissipates as the climate warms, to the point that there is almost no observable seasonality for the higher-concentration scenario during the last two decades of this century. This finding suggests there is the potential for the dengue season to extend all year, with outbreaks occurring at any time. Consequently, disease surveillance and control activities would need to be geographically and temporally adapted to mitigate dengue epidemic risk in response to climate change.

## Introduction

Dengue is an acute viral disease which has spread around the world causing epidemics and establishing itself in endemic transmission cycles. Dengue virus infections are the most common vector-borne human disease, accounting for an estimated 105 million infections globally per year, of which 51 million manifest the disease at some level of clinical or sub-clinical severity [1]. The virus is transmitted to humans by infected female mosquitoes of the *Aedes* species, most commonly *Ae aegypti*, which is associated with major epidemics, and to a lesser extent by *Ae albopictus*, which is a less efficient vector [2, 3]. Infection with dengue virus is often asymptomatic but can result in a wide spectrum of disease manifestation, from undifferentiated fever to potentially life-threatening dengue shock syndrome and dengue hemorrhagic fever [4].

Bangladesh, a densely populated country, is one of the Asian countries most affected by dengue epidemics in recent years. In 2000, the first reported outbreak of dengue occurred with at least 5,551 cases and 93 deaths recorded across three major cities [5]. Since then, dengue cases and deaths have been reported each year, mostly during monsoon and post-monsoon season, with some year-to-year variation [6]. The reported number of dengue cases and associated deaths is estimated to be considerably lower than actual [7]. Through a nationwide serosurvey it was estimated that by the end of 2015, 40 million people (95% CI: 34-47) had been infected with dengue virus at some point during their lives with 2.4 million (95% CI: 1.3-4.5) new infections occurring each year [8].

Dengue incidence is much higher in the three major urban conurbations of Dhaka, Chittagong, and Khulna, where ∼50% of the urban population live, in comparison to the more dispersed rural villages in the rest of the country [8]. Dengue is a highly climate-sensitive disease because the vector, *Aedes* mosquitoes, is profoundly influenced by changes in mean temperature and temperature variations [9]. Due to climate change, the annual mean temperature of Bangladesh increased at an average rate of 0.02°C per year between 1971 and 2010, which is considerably higher than the global average increase [10]. Projected future increases in mean temperature for Bangladesh relative to pre-industrial period (1861-1880) are as much as 3.2 to 5.8°C by the end of the 21^st^ Century for a high greenhouse gas emission scenario [11]. In this context, understanding the relationship between dengue incidence and climate variables is crucial to understand the future burden of dengue in Bangladesh and to inform appropriate public health responses.

Previous studies have sought to understand aspects of dengue transmission that depend on temperature and these, in turn, have informed the evaluation of the potential effect of climate change on the future spread of dengue infections. One approach is to calculate the vectorial capacity (VC), which is used to describe the epidemic potential of a vector-borne disease and is closely related to the risk of outbreaks occurring [12]. Vectorial capacity was originally derived from a malaria transmission model [13]. It is related to the basic reproduction number and hence provides a threshold for an epidemic to occur [14, 15]. Recent formulations of VC now include temperature dependent parameters that characterise host, virus, vector, and interaction factors related to transmission [14].

Temperature observations obtained from weather stations have been used to identify historical trends in VC and, thus, to explain past dengue distribution [16]. Quantifying potential future changes in VC requires information about the future climate. Global Climate Models (GCMs) are complex mathematical representations of the major climate system components (atmosphere, land surface, ocean, and sea ice) and their interactions and are one source of information about future climate conditions. Although GCM simulations are imperfect representations of the climate, especially at regional scales, and do not represent all potential scenarios for future climate change, information derived from them can be used to generate scenarios to support the exploration of climate change impacts.

The Coupled Model Intercomparison Project phase 5 (CMIP5) [17] provides simulations of many different GCMs of the Representative Concentration Pathway (RCP) scenarios specified by the Intergovernmental Panel on Climate Change (IPCC) [18]. The RCPs provide information on possible future trajectories for the main forcing agents of climate change, for example, atmospheric concentrations of greenhouse gases and other air pollutants and changes in land use [18]. Varying underlying assumptions and simulation approaches mean that the different CMIP5 GCMs simulate varying future climate projections for the same RCP. Assessments of future climate change or its impacts usually address this uncertainty by using multiple GCM outputs for analysis [19]. Further, the outputs for GCMs can be biased relative to climate observations. For example, a GCM can be cold- or warm-biased relative to temperature observations. One contributing factor when observations for specific sites or small areas are considered is that GCM output lacks spatial detail and typically represents average climate conditions across 100s of kilometers. When analysing climate impacts for a specific region, the output from GCMs needs to be downscaled and bias-adjusted. This is particularly important when assessing variables that respond in a non-linear manner to changes in climate variables [20], as is the case with VC.

Earlier studies projecting the impact of climate change on dengue focused on global or broad regional scales and made use of GCM output for different future climate change scenarios at a global or regional level. For example, Liu-Helmersson *et al*. (2016) investigated historic and projected dengue epidemic potential in Europe for different climate change scenarios using VC calculations [16]. We performed a country-specific study for Bangladesh using bias-adjusted data from multiple GCMs to provide a range of climate scenarios for the 21^st^ Century. In response to these scenarios, we investigated the projected changes in VC.

## Methods

### Collection of climate data

#### Observed temperature data for Bangladesh

The Bangladesh Meteorological Department measures weather parameters at 35 stations that are well-distributed across the country (Figure 1). We received daily temperature observations (mean, minimum and maximum) at these weather stations for the period 1975-2016 [21]. We checked the temperature values in the downloaded dataset by comparing to the data reported in published literature and searching for extreme values (Figure S1 in the supplementary material).

**Figure 1:**
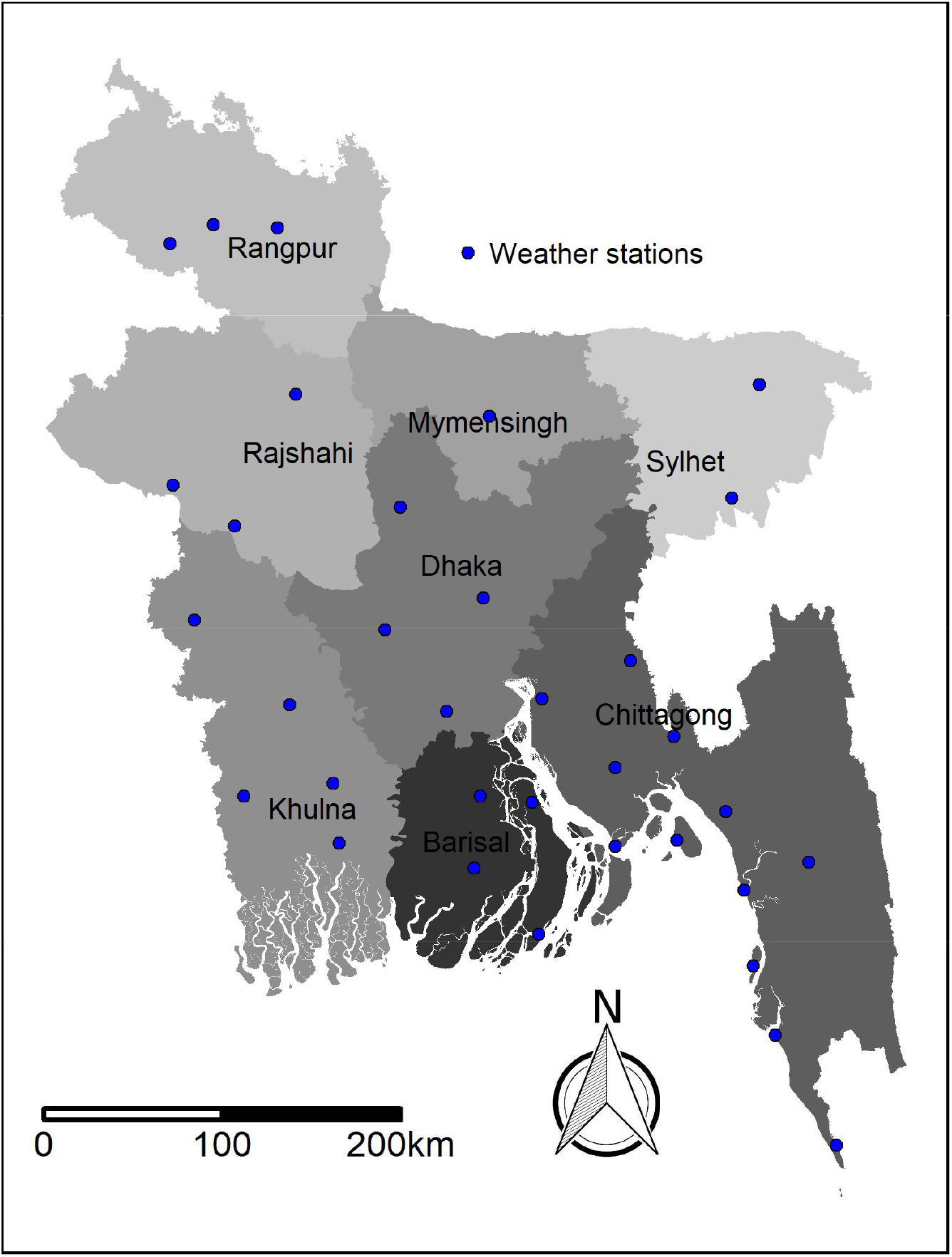
Map of Bangladesh showing divisional boundaries and the location of the 35 weather observation stations across the country (blue dots).

#### Climate data from Global Climate Models

Bias-adjusted daily temperature data derived by the Inter-Sectoral Impact Model Intercomparison Project (ISI-MIP) from five CMIP5 GCMs (HadGEM2-ES, GFDL-ESM2, IPSL-CM5A-LR, MIROC-ESM-CHEM and NorESM1-M) have been used previously for climate change impact assessments, projecting dengue epidemic potential in Europe, for example [16]. ISI-MIP applied a statistical bias-adjustment approach to correcting GCM data towards an observed reference climate dataset for the 1960-1999 period (the WATCH Forcing Data) [22]. More recently, the ISIMIP2b project applied the same bias-adjustment method to a different set of GCMs (HadGEM2-ES, GFDL-ESM2, IPSL-CM5A-LR, MIROC5) towards a newly compiled reference dataset EWEMBI [23, 24]. NorESM1-M was disregarded because of a lack of wind data and, pertinently to the climate of Bangladesh, MIROC-ESM-CHEM was replaced by MIROC5 due to a better representation of the monsoon by the latter [24]. For each of the four GCMs, daily gridded (0.5° x 0.5°, approximately 50 km^2^, resolution) temperature data were available for a single historical dataset for the period 1950-2005 and for four different datasets corresponding to four RCP scenarios for the period 2006-2099 [25].

Of the four RCP scenarios, we evaluated RCP 4.5, which corresponds to a greenhouse gas emissions pathway that peaks in 2040 and then declines to 1960s emission levels by 2090, and RCP 8.5, which corresponds to emissions that continue to increase significantly until near the end of the 21^st^ Century. RCP 4.5 was selected because it is an ambitious pathway developed from United Nation’s 2015 Paris Agreement to reduce the chance of significant disruption to the Earth’s natural cycles [26]; and RCP 8.5 was selected because it broadly reflects a scenario that best matches recent emissions and a future in which little additional action on reducing greenhouse gas emissions is taken.

To compare temperature data derived by ISI-MIP with observed temperature data in Bangladesh, daily temperature data for the 35 weather station locations shown in Figure 1 were extracted from gridded ISI-MIP data using Climate Data Operators (version 1.9.8), using a collection of command line operators to manipulate and analyze climate model data [27]. Annual cycles of monthly averaged mean, maximum, and minimum temperature for 1986-2005 for the observed and ISI-MIP datasets at all 35 weather stations were compared through visual inspection. Good alignment of annual temperature cycles was observed for all weather stations except Dinajpur, where a warm bias was observed in daily maximum temperature (Figures S2, S3, and S4; details in Supplementary Material).

### Calculation of Vectorial Capacity

To calculate VC, we used the approach derived in Liu-Helmersson *et al*. (2016) [16]. This approach and similar formulations have been used in several studies to link changes in temperature to the potential for dengue transmission [12, 15, 28]. VC is given by:

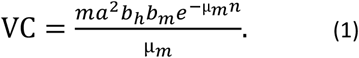

Equation 1 has six vector-related parameters that depend on temperature: the average vector biting rate (*a*); the probability of vector-to-human transmission per bite (*b*_*h*_); the probability of human-to-vector infection per bite (*b*_*m*_); the duration of the extrinsic incubation period (*n*); the vector mortality rate (*μ*_*m*_); and the female vector-to-human population ratio (*m*). The temperature relationship of each of the vector parameters was derived from previously published literature for the primary vector *Ae. aegypti* [15] and are included in the Supplementary Material (Table S1). Assuming the infectious period for dengue is 5 days, then when VC reaches a threshold value of 0.2 per day, one infected person will infect at least one other person in a dengue naïve population while infectious [15, 29].

VC varies from hour-to-hour throughout the day with temperature. However, hourly temperature data are not available from many observation and GCM-derived climate datasets. We sourced daily values of mean, maximum and minimum temperature from climate data sources. To account for the impact of hour-to-hour temperature fluctuations within each day, a sinusoidal hourly temperature variation between maximum and minimum temperature, diurnal temperature range (DTR), within a period of 24 hours was assumed [15]. Vectorial capacity was calculated for hourly temperature points and then averaged to yield a daily mean estimate. Daily VC at eight divisions, the first level administrative unit in Bangladesh, were calculated by averaging the daily VC of the weather stations located within divisional boundaries to facilitate regional analysis. There are four distinct seasons in Bangladesh: the dry winter season (December to February); the pre-monsoon hot summer season (March to May); the rainy monsoon season (June to September); and the post-monsoon autumn season (October to November). From the daily VC we then calculated monthly, seasonal, and annual VC for each GCM at the divisional level and the overall national level.

Our results and figures were produced using R version 4.0.2 [30] with the packages ‘ncdf4’ (extraction of temperature data from climate model netCDF files) [31] and ‘tidyverse’ (data cleaning, summarisation and plotting) [32].

## Results

### Annual average vectorial capacity

The national average annual VC of *Ae. aegypti* mosquitoes in Bangladesh over 1975-2016 ranged between 0.7 and 1.1 per day when calculated using the observed data from weather stations. This is well above the threshold value for epidemic dengue transmission (0.2 per day) and reflects the ongoing dengue outbreaks that have occurred since 2000. The national average annual VC declined slowly but significantly at an average rate of 0.002 per year over the period 1975 to 2016 (p < 0.001) (Figure S5 in the supplementary material). Some regional differences over the same period were observed with lowest average annual VC in Sylhet (0.8, range: 0.8 – 0.9) and highest average annual VC in Barisal (1, range: 0.9-1.1).

We validated the VC results from the ISI-MIP data by comparing them with the observationally derived VC data over the 1986-2005 period. Regional annual VC calculated with the four ISI-MIP GCMs for 1986-2005 period fluctuated around the 20-year (1986-2005) average of regional annual VC calculated with observed data (Figure 2). This indicates that the annual VC values calculated with the ISI-MIP data were comparable to the annual VC calculated with observed temperature data, giving us some confidence in future projections of VC derived from the ISI-MIP data.

**Figure 2:**
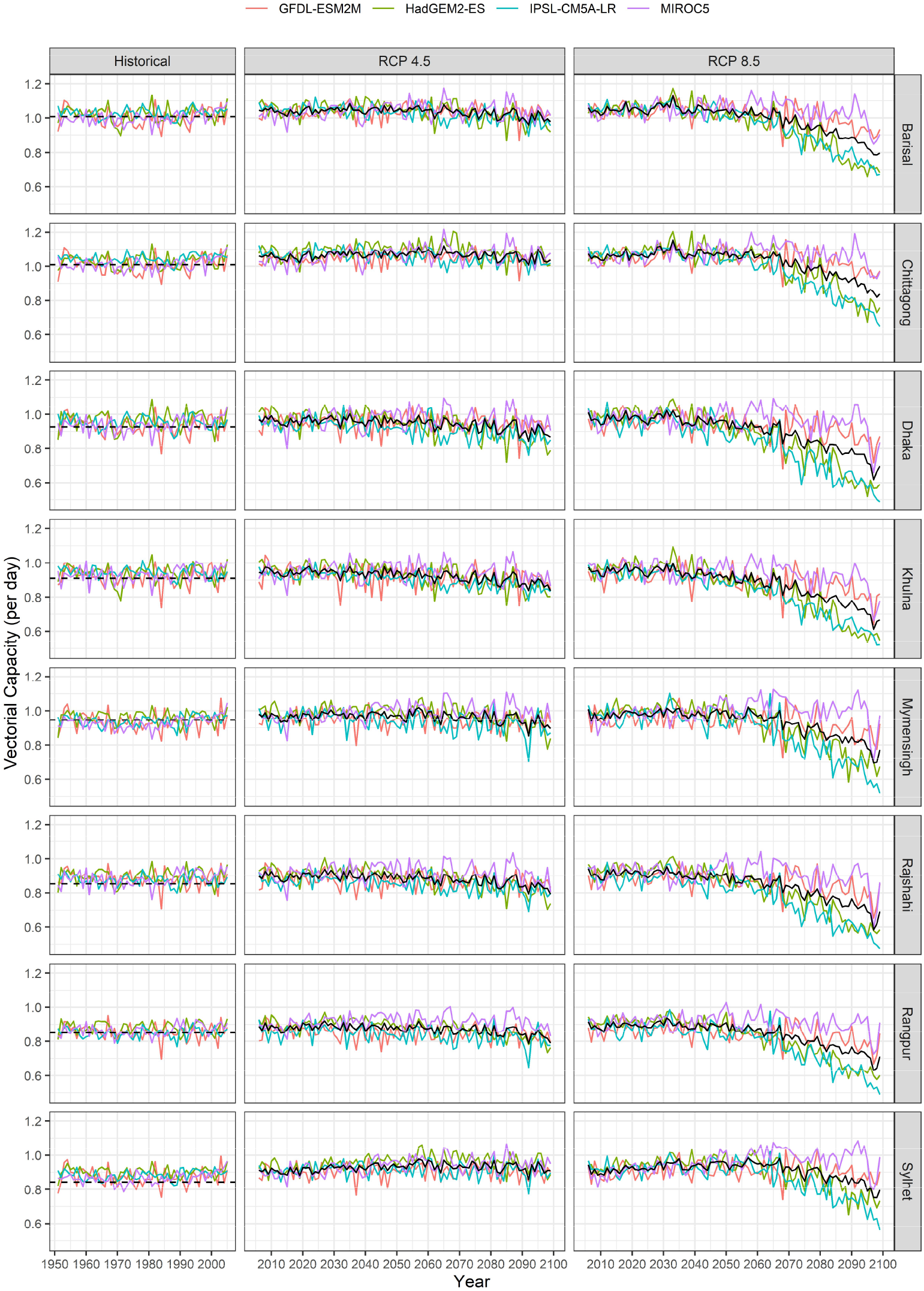
Average annual vectorial capacity of *Ae. aegypti* mosquito in Bangladesh over 1951-2099 for each division of Bangladesh and each ISIMIP GCM under RCP4.5 and RCP8.5. The dashed black line represents average annual vectorial capacity over 1986-2005 calculated with observed data. The solid black line represents results averaged over the four GCM.

We now consider future changes in average annual VC projected by the ISI-MIP data. For all four GCMs, the average annual VC in 2030-49 period for the lower-concentration scenario, RCP 4.5, was found to increase slightly in Chittagong and Sylhet and to remain unchanged in Dhaka and Khulna, relative to the 1986-2005 average annual VC (Figures 2 and 3). For the same RCP scenario, a slight decrease in the average annual VC was noted in Dhaka, Khulna, and Rajshahi divisions over the last two decades of the 21^st^ Century, relative to 1986-2005, particularly for the HadGEM2-ES and IPSL-CM5ALR models. For RCP 8.5, average annual VC over the 2030-2049 period was found to increase in Chittagong and Sylhet divisions for all four GCMs compared to 1986-2005. A pronounced decline in VC begins in 2060 for RCP 8.5 in all divisions and continues for the remainder of the century for all four GCMs. The magnitude of the decline differs between GCMs, being most pronounced for HadGEM2-ES and IPSL-CM5A-LR and divisions, being most pronounced for Dhaka and Khulna (Figures 2 and 3).

**Figure 3:**
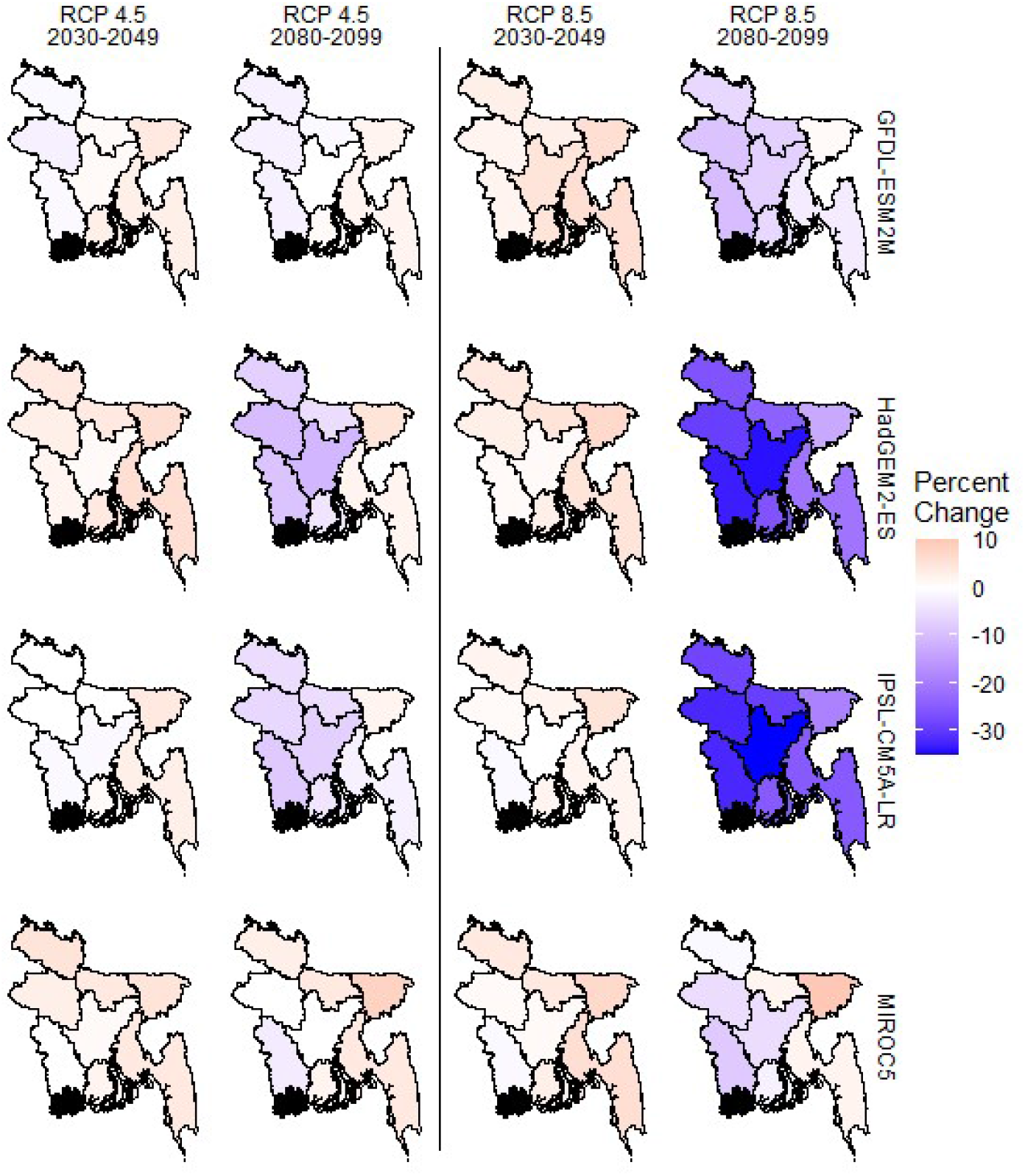
The spatial distribution of average vectorial capacity in each division of Bangladesh for two twenty-year periods for the four ISIMIP GCMs under RCP4.5 and RCP8.5. The change of vectorial capacity for each model is relative to its own vectorial capacity for 1986-2005.

Irrespective of RCP, the annual VC calculated for all four ISI-MIP GCMs for all years in the 1950-2099 period remained above 0.2 in all eight divisions (Figure 2). This indicates a level suitable for potential dengue epidemic transmission at least some time during all of these years.

### Seasonality of vectorial capacity

Strong seasonality of VC was found with the observed data, with VC peaking during the rainy monsoon season and reaching a minimum during the dry winter season. Similar seasonality patterns of VC for all four ISI-MIP models were found for the historical data (1986-2005) as well as for the two RCP scenarios for the 2030-2049 period in all eight divisions (Figure 4). The seasonal cycle in VC for all four ISI-MIP models became less pronounced during 2080-2099 for RCP 4.5, with increased VC during the winter/dry months and decreased VC during the monsoon months. During the last two decades of the 21^st^ Century, the four ISI-MIP models differ in their results for RCP 8.5. There is relatively little seasonality in VC for the GFDL-ESM2M and MIROC5 models, though it is still generally highest during the monsoon. However, the seasonal cycle is inverted for the HadGEM2-ES and IPSL-CM5A-LR models, with lower VC during the monsoon compared with the winter/dry months (Figure 4).

**Figure 4:**
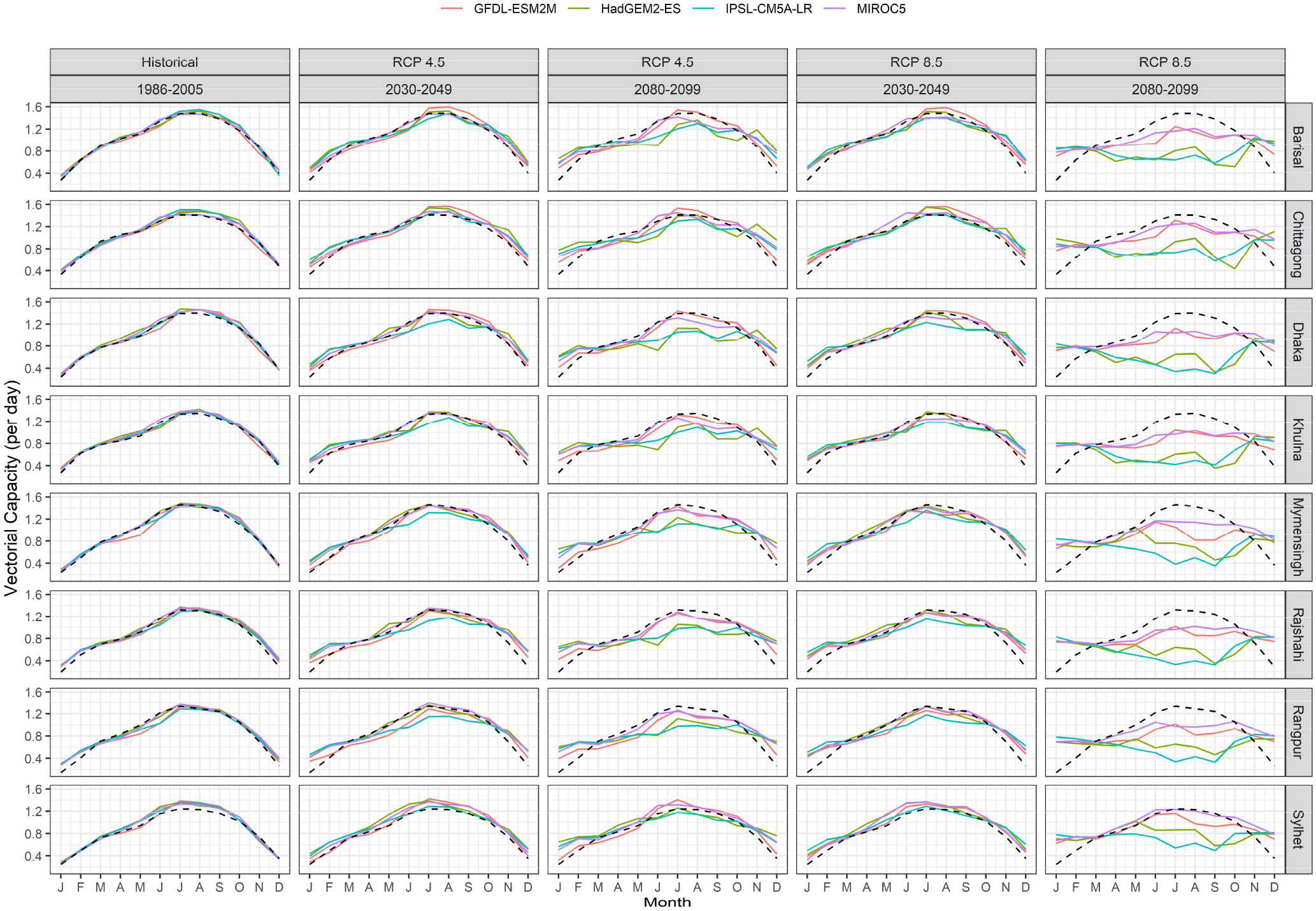
Monthly average vectorial capacity of *Ae. aegypti* mosquitoes for three twenty-year periods in each division of Bangladesh for each ISIMIP GCM under RCP4.5 and RCP8.5. The dashed line represents monthly average vectorial capacity calculated with observed data, 1986-2005.

Although VC is a function of both mean temperature and DTR (Figure 5), these seasonal differences in VC are driven by seasonal changes in mean temperature. The optimal temperature for *Ae. aegypti* to transmit dengue is around 29.3°C when DTR=0°C. The climate change scenarios for both RCPs and all four GCMs have little change in DTR. However, the RCP 8.5 scenario of all four GCMs is consistent with increasing mean temperature during both monsoon (i.e., away from the optimum temperature) and winter (i.e., towards the optimum temperature). These changes are more prominent towards the end of the century and with the HadGEM2-ES and IPSL-CM5A-LR models. The RCP 4.5 results are due to this mechanism as well, only less pronounced due to smaller warming.

**Figure 5:**
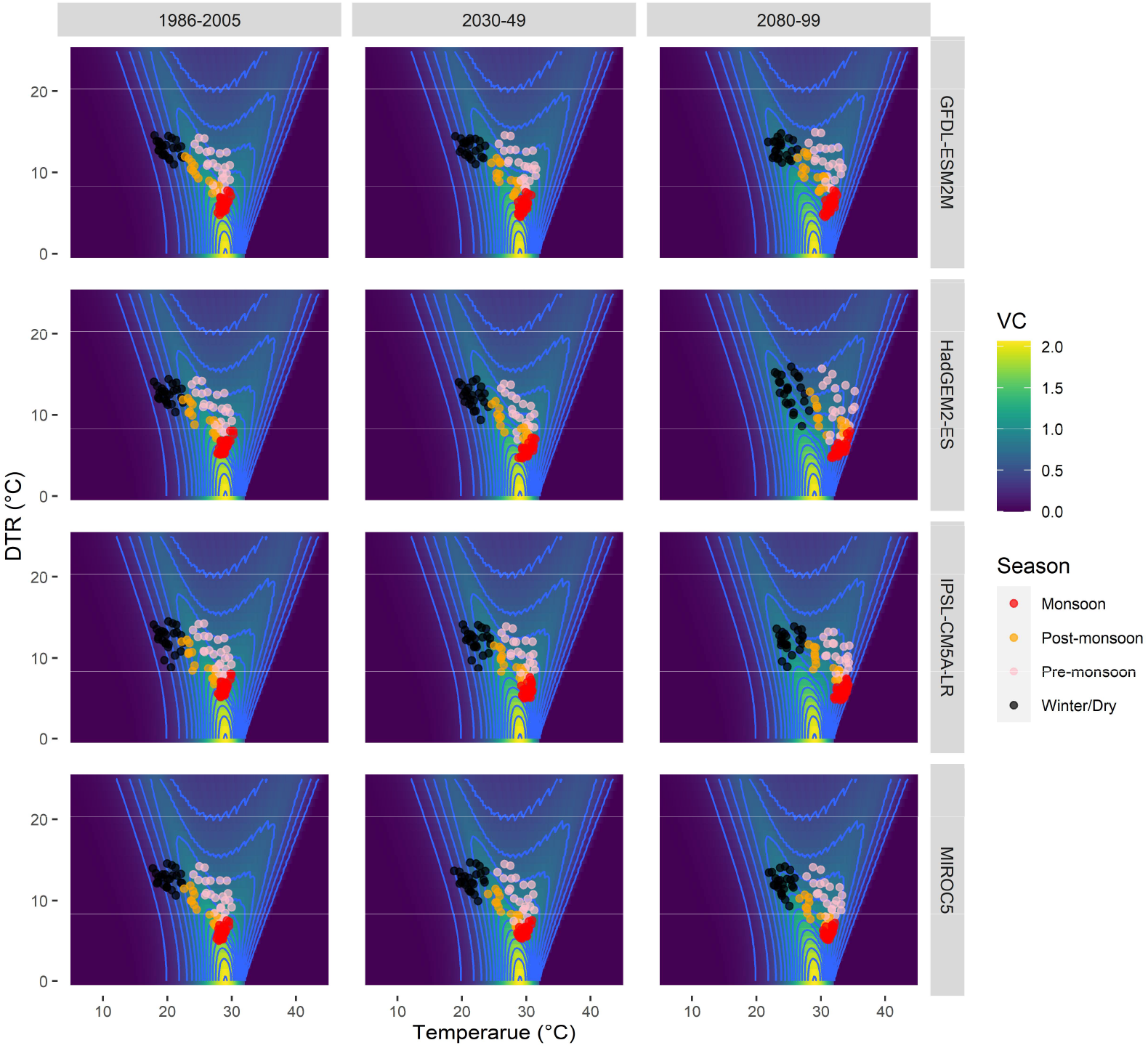
Vectoral capacity as a function of temperature and diurnal temperature range. Seasonal temperature and diurnal temperature data for three twenty-year periods for RCP8.5 from the four ISIMIP GCMs are superimposed.

Irrespective of RCP, the monthly average VC remains well above the threshold value of 0.2 per day throughout the year for all four ISIMIP GCMs for the entire 21^st^ Century, indicating potential for epidemic dengue transmission at all times of the year.

## Discussion

We estimated the VC for *Aedes aegypti* mosquitoes to explore the potential of dengue transmission at a sub-national level for Bangladesh to the end of the 21^st^ Century. Our analysis showed that annual VC for all divisions in Bangladesh is projected to be substantially above the threshold for epidemic dengue transmission throughout this century. During the second half of the century, the annual VC is projected to decline compared to the current level, particularly for the RCP 8.5 scenario. However, the fall in annual VC masks substantial changes in monthly VC, with the VC for the winter/dry season increasing to a level close to a reduced monsoon peak. This suggests that in the future, the dengue fever season could become longer, with outbreaks occurring at any time of the year.

Projections of VC may provide insight as to where and when there is likely to be a risk of transmission given the existence of other relevant factors [13]. The decline in VC for RCP 8.5 for all four GCM simulations during second half of the 21^st^ Century is consistent with continued increase in global greenhouse gas emissions resulting in an increase in temperature to sub-optimal conditions for the mosquito vector. This long-term projection of decreased dengue risk is in contrast with earlier projections for Bangladesh, where a rise in dengue case numbers was predicted with an assumed increase in temperature of 3.3°C by the end of the 21^st^ Century [33]. The contrasting directions of these two projections might be explained by the fact that in the earlier study, a positive linear relationship of notified dengue cases and temperature was assumed, whereas the relationship of temperature and vector parameters of VC is non-linear with high temperatures being sub-optimal for *Aedes aegypti* mosquito behaviour and survival (Figure 5) [15]. The regional differences in annual VC change to the mid-21^st^ Century highlight the need for improved disease control activity in eastern and north-eastern Bangladesh over the coming decades.

Vectorial capacity shows strong seasonality in both temperate and tropical regions [16]. For the recent past, current, and projected near future climate of Bangladesh, strong seasonality of VC is similarly observed in this study. Dengue epidemic potential projected for the European cities by Liu-Helmersson *et al*. (2016) [16] showed that VC is only sufficiently high for dengue transmission to occur for a few months in a year. This is explained by the strong seasonal cycle in temperature in European cities. However, as in other tropical cities (e.g., Singapore, Colombo, and Miami) [16], VC in Bangladesh never drops below the transmission threshold (0.2 per day), indicating year-round potential for transmission. Despite the potential, almost no dengue cases were recorded in past two decades during winter/dry season in Bangladesh due to lack of rainfall required for replenishment of common mosquito breeding sites [7, 34]. However, during the latter part of the 21^st^ Century, projected seasonality is quite different from what we experience now, particularly for the RCP 8.5 scenario. Season to season variability is expected to decline due to an increase in temperature above the optimal in the monsoon period and towards the optimal during winter, though the projected change does not lead to a decline in VC below the 0.2 threshold value.

Since GCMs are not perfect representations of the climate system, it is important to consider the robustness of these results to the reliability of the GCM information used. Although we have used bias adjusted GCM data, we recognise that bias adjustment may not resolve issues with the simulation of climate processes that affect simulated future climate changes. We note previous work that has highlighted deficiencies in the IPSL-CM5A-LR and MIROC5 models [36]. However, even if these models were disregarded, our conclusions would be unchanged as the GFDL-ESM2M and HadGEM2-ES models also support these conclusions. We also note that the set of GCMs that we have used spans the range of temperature increase simulated for the region by the full range of CMIP5 GCMs [37]. This means that we have sampled the uncertainty in future VC changes related to uncertainty in future warming simulated by CMIP5.

It is also worth considering the robustness of our results to the parameter values used in our calculation of VC. The point estimates of vector parameters used in this study were obtained from fitted experimental data on *Ae. aegypti* in different regions of the world, incorporating uncertainties around these and specific empirical estimates for the region would improve our VC estimate. However, for the range of temperature used in this analysis, the optimum VC temperature is unlikely to be very different for different parameter values [15], giving us confidence that our conclusions are robust to the VC parameter values.

Finally, we did not consider other changes in climatic variables such as rainfall and humidity, which could be important drivers of dengue risk [33], especially at a seasonal or regional level. For example, currently, the winter is dry in Bangladesh and hence, there are fewer mosquito breeding sites and outbreaks of dengue do not occur despite VC being above 0.2 [35]. Higher rainfall is projected for this season by regional climate models which means this risk [11] could be realised or further increase dengue outbreak potential in this season, relative to our results for the late 21^st^ Century.

As long as the dengue virus, mosquito vectors, susceptible population and suitable environmental conditions prevail, there is no reason to curtail ongoing prevention strategies or delay adoption of newer and more effective measures for disease control. Possible seasonal and geographic extension of dengue risk throughout latter half of this century suggests that the spatiotemporal coverage of dengue surveillance and vector control activities will need to be broadened to mitigate future epidemics of dengue in Bangladesh.

## Acknowledgements

We want to thank the Bangladesh Meteorology Department for providing daily weather data, and modeling teams of ISI-MIP and DECCMA for providing climate model output used in this study.

## Data availability statement

All data and calculation/analysis code are publicly available from https://github.com/The-Kirby-Institute/VC_changing_climate_BD.

